# TMPRSS2 activation of Omicron lineage Spike glycoprotein is regulated by Collectrin-like domain of ACE2

**DOI:** 10.1101/2023.09.22.558930

**Authors:** Anupriya Aggarwal, Christina Fichter, Vanessa Milogiannakis, Timothy Ison, Mariana Ruiz Silva, Camille Esneau, Nathan Bartlett, Louise M Burrell, Sheila K Patel, Melissa Churchill, Thomas Angelovich, Greg Walker, William Rawlinson, Malinna Yeang, Jen Kok, Vitali Sintchenkov, Deborah Chandra, Andrea Kindinger, Anouschka Akerman, Rhys Parry, Julian D Sng, Greg Neely, Cesar Moreno, Lipin Loo, Anthony D Kelleher, Fabienne Brilot, Alexander Khromykh, Stuart G Turville

## Abstract

Continued high-level spread of SARS-CoV-2 has enabled an accumulation of changes within the Spike glycoprotein, leading to resistance to neutralising antibodies and concomitant changes to entry requirements that increased viral transmission fitness. Herein, we demonstrate a significant change in angiotensin-converting enzyme 2 (ACE2) and transmembrane serine protease 2 (TMPRSS2) dependent entry by primary SARS-CoV-2 isolates that occurred upon arrival of Omicron lineages. Mechanistically we show this shift to be a function of two distinct ACE2 pools based on TMPRS22 association with the ACE2 Collectrin-Like Domain (CLD). In engineered cells overexpressing ACE2 and TMPRSS2, ACE2/TMPRSS2 complexes led to either augmentation or attenuation of viral infectivity of pre-Omicron and Omicron lineages, respectively. Mutagenesis of the ACE2-CLD TMPRSS2 cleavage site in ACE2 restored infectivity across all Omicron lineages through enabling ACE2 binding that facilitated TMPRSS2 activation of viral fusion. Our data supports the evolution of Omicron lineages towards the use of ACE2 unable to form complexes with TMPRSS2 and consistent with ACE2 structure and function as a chaperone for many tissue specific amino acid transport proteins.

**Graphical Abstract:**
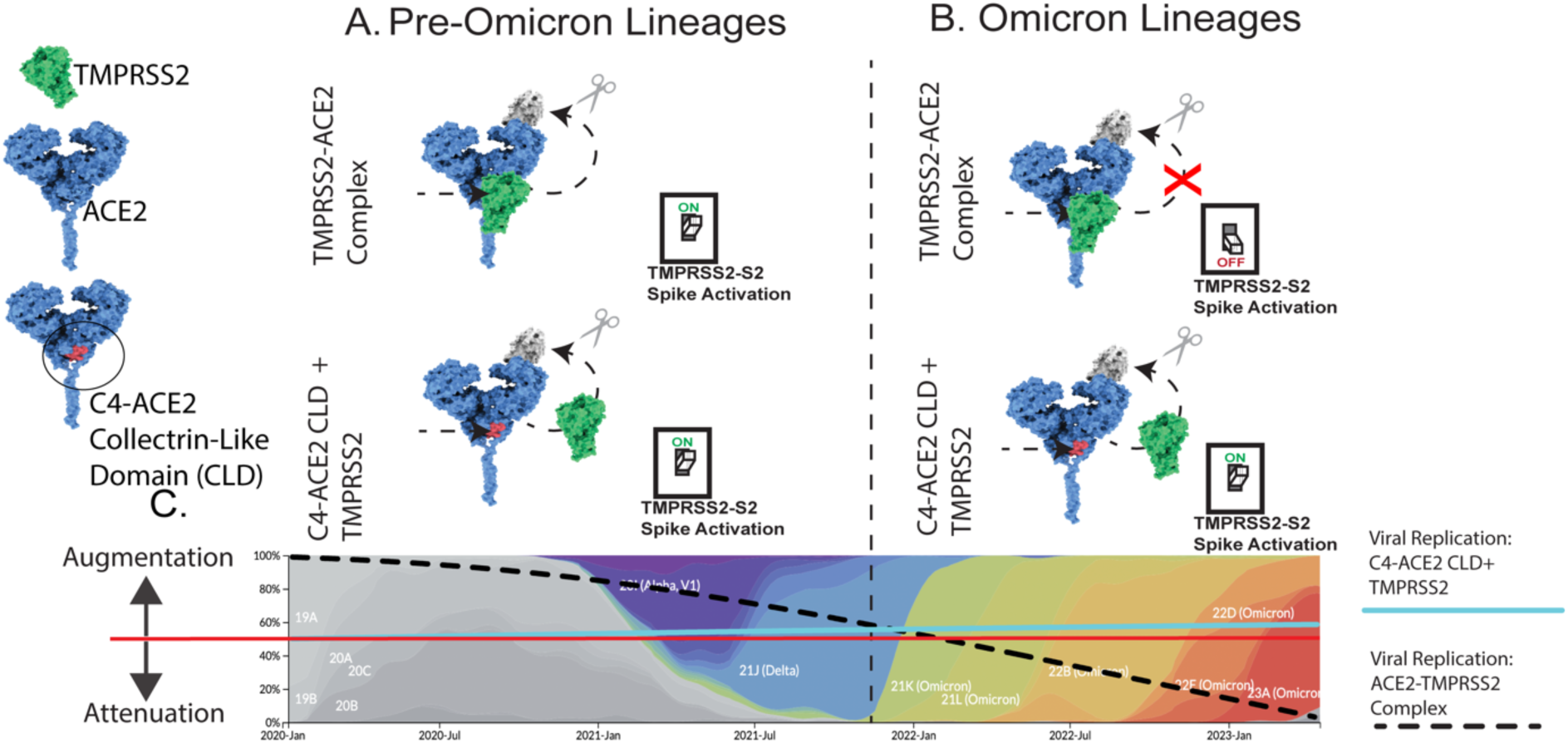
ACE2-TMPRSS2 pool model and evolution of SARS-CoV-2 tropism. **A. &-B. ACE2-TMPRSS2** pool model to reconcile the evolving entry requirements of SARS-CoV-2 and changes in viral tropism *in vivo*. **A.** Both SARS-CoV-1 and early SARS-CoV-2 (pre-Omicron) lineages have molecular dual tropism, with efficient entry when ACE2 (blue protein) can form complexes with TMPRSS2 (green protein) and in settings where TMPRSS2 is excluded from ACE2 (C4-ACE2 CLD). In both settings ACE2 initially engages SARS-CoV-2 spike and fusion is then triggered through TMPRSS2 cleavage of the Spike S2 domain **B.** Over time, the dual tropism for two distinct pools of ACE2 (with and without TMPRSS2) has been lost, with consolidation towards ACE2 where TMPRSS2 is no longer in a complex. With the arrival of Omicron lineages, ACE2-TMPRSS2 complexes could no longer enable efficient Spike S2 cleavage and fusogenic activation (TMPRSS2 “off” confirmation”). Rather, only TMPRSS2 uncoupled from ACE2 could facilitate the latter cleavage of S2. C. Over time this has further consolidated over generations of omicron lineages from 2022 lineages (BA.1, BA.2 and BA.5) through to 2023 lineages (XBB.1.5) and now in 2024 JN.1 lineages such as KP.3. Overall, this supports the initial molecular tropism of early SARS-CoV-2 clades to be similar to that observed for SARS-CoV-1, with dual tropism across both ACE2 pools and replication proceeding in tissues where ACE2-TMPRSS2 complexes would be prevalent (e.g. Lung). The evolution away from ACE2-TMPRSS2 complexes towards ACE2 where TMPRSS2 is structurally uncoupled (e.g. ACE2 as a chaperone for solute carriers SLCA619 or SLCA620) is consistent selection of this ACE2 pool in a manner that has sustained transmission fitness within the human population.

## Introduction

Members of the *Coronaviridae* family readily circulate within the human population with four common human coronaviruses (HCoVs; 229E, OC43, NL63, and HKU1) known to cause mild to moderate upper respiratory tract infections. In contrast, severe acute respiratory syndrome coronavirus (SARS-CoV-1), Middle East respiratory syndrome (MERS-CoV) and SARS-CoV-2 are associated with a greater risk of severe respiratory disease. Whilst the manifestations of clinical disease are complex and influenced by prior vaccine and convalescent immunity, a greater understanding of cellular entry pathways and viral tropism in coronaviruses is required to resolve the viral pathogenesis that eventually leads to clinical disease. Furthermore, many viruses, as they evolve, change how they target cells in different tissues, causing organ-specific disease and outcomes. Early clades of SARS-CoV-2 had an entry pathway aligned with that of SARS-CoV-1, where ACE2 was the primary receptor and fusion was activated by protease cleavage. The serine protease TMPRSS2 facilitated this at the membrane and the cysteine protease Cathepsin L within endosomes. Whilst the latter pathway is seen as a redundant mechanism of entry for the virus *in vitro*, murine knockout studies and *in vitro* infections, highlight the virus’s preference for membrane entry TMPRSS2^2^ mediated fusion. However, transmission and infection *in vivo* can take place across many tissues, with efficient entry into one tissue type often differing from another ^3–6^.

Early variants of concern (VOC) in the COVID-19 pandemic followed a similar trajectory in tropism, which involved engagement of ACE2 with Spike subunit 1 (S1) domain and then fusogenic activation through cleavage of Spike subunit 2 (S2) by TMPRSS2 ^7,8^. ^9,10^. In brief, fusogenic activation of Spike requires proteolytic cleavage at two distinct sites, S1/S2 and S2. Cleavage of S1 and S2 is mediated by the proprotein convertase Furin during biosynthesis of Spike within infected cells, leading to cleaved S1- and S2 subunits that remain non-covalently linked. The S1 subunit contains the receptor binding domain, which binds ACE2, whilst S2 harbours the TMPRSS2 cleavage site. This is located immediately upstream of the hydrophobic fusion peptide, which promotes viral membrane fusion. In a cell type-dependent manner, prior cleavage of S1/S2 by Furin increases the ability of S2 to be cleaved by TMPRSS2, which culminates in greater levels of viral fusion. The first generation of VOCs (Alpha, Beta and Gamma) demonstrated a fitness gain that was primarily mediated by increases of ACE2 affinity through Spike polymorphisms, such as N501Y ^9,10^. The initial dominance of the VOC Alpha and subsequent appearance of the VOC Delta resulted from key changes (P681H and P681R) at the Furin cleavage site within Spike. These changes led to fitness gains primarily through efficient TMPRSS2 cleavage of S2. As a result, cell types with relatively high levels of TMPRSS2, including those from the lower respiratory tract, have augmented permissiveness to variants with efficient Furin-mediated Spike cleavage, such as Delta^11–15^.

The emergence of BA.1 and BA.2 Omicron lineages shortly after Delta was a seismic shift at two levels. Firstly, the ability of Omicron lineages to efficiently evade neutralizing antibodies, and secondly, a shift in tropism away from the lower respiratory tract ^3–5,14–16^. This combination has driven global infection waves that surpassed those prior to Delta. Whilst Omicron lineages have further diversified and in some cases recombined to create greater diversity, the fundamental understanding of the entry requirements of Omicron SARS-CoV-2 remains unclear at two levels. Firstly, the mechanistic basis of the tropic shift observed upon the arrival of Omicron BA.1/BA.2. Secondly, as the virus continues to evolve, there is uncertainty as to whether the virus is maintaining an Omicron entry pathway trajectory or, alternatively, regressing to the entry pathway observed during and prior to Delta. Herein, using genetically intact clinical isolates that span the pandemic to date, we observed viral replication across a continuum of engineered cells to resolve the entry requirements and/or restrictions of each variant. As a result, we have defined the Omicron tropic shift to be a consequence of the ability of ACE2 to form complexes with TMPRSS2. The latter ACE2-TMPRSS2 complex is physiologically relevant within the renin angiotensin aldosterone system (RAAS) pathway, as it enables regulation of ACE2 shedding by a combination of exclusion of ADAM-17 and/or through proteolytic cleavage of ACE2 at its CLD, that in turn removes increases its endosomal uptake ^17^. For pre-Omicron and Omicron lineages, the presence of the ACE2-TMPRSS2 complex augments or attenuates TMPRSS2 Spike activation, respectively. This provides the mechanistic basis of the change in viral tropism upon the arrival of Omicron and reconciles discrepant *in vitro* and *in vivo* observations of TMPRSS2 Spike activation. It further highlights, SARS-CoV-2’s promiscuous use of ACE2 pools shortly following zoonosis in a manner that is similar in mechanism to SARS-CoV-1.

## Results

### Permissiveness of cell lines expressing ACE2 and TMPRSS2 to SARS CoV-2 variants that have dominated throughout the pandemic

Throughout the pandemic, we have collaborated with genomics surveillance units to enable real-time isolation and culture of primary SARS-CoV-2 isolates. This was enabled by cellular substrates that were genetically engineered cell lines to increase permissivity to SARS-CoV-2 ^16,18–20^ Through increasing expression of ACE2 and TMPRSS2 (both co-expressed in known targets of the respiratory and gastrointestinal tract^7,21–24^), both ourselves and others attempted to emulated the viral entry pathway observed in primary cells and *in vivo*. The HEK293T and VeroE6 cell lines have been extensively used in virological assays as starting substrates, as they lack key host restriction mechanisms and as such are suited for assay platform development and viral propagation ^16,18–20^. Whilst the latter approach was used successfully pre-Omicron, evidence rapidly emerged demonstrating that Omicron lineages no longer effectively utilized the ACE2-TMPRSS2 pathway in various cell lines and primary cell settings ^14,16,25–27^. To increase the efficiency of primary isolations from nasopharyngeal swabs, we screened existing cell lines for susceptibility to infection across viral variants representative of the major variants globally from the last three years (Fig. 1A). We utilized a high throughput imaging approach with live cell nucleus staining to enumerate the accumulation of viral cytopathic effects through the dose-dependent loss of nuclei ^16,20^ (Fig. 1B). Combined with automated nuclei counting algorithms, this provided a non-biased method of screening variant panels against various cell lines. Using this approach, we observed a continuum of viral permissiveness in each cell line (Fig. 1C-G). As expected, the VeroE6 line was the least permissive in contrast to two VeroE6 lines and a ACE2-Hek 293T line engineered to express TMPRSS2 (HAT-24) (Fig. 1C-G). Expression of TMPRSS2 in the same parental VeroE6 line or the HAT-24 line did not lead to equivalent outcomes in infection with each lineage. The VeroE6-T2 cell clone provided the greatest breadth for viral replication across all lineages (Fig. 1G) with titers ten-fold higher than the VeroE6-T1 cell clone for all lineages with the exception of BA.2 and BA.2 (Fig. 1F versus 1G). Curiously, the HAT-24 line provided a similar profile of lineage permissiveness to the VeroE6-T1 line (Fig.1H), with the exception of higher replication observed in lineages BQ.1.2 and XBB.1.5. Given the unique breadth across variants with the VeroE6-T2 line, we expanded all viral isolates over 24 hours using this clone and observed high RNA copies/ml of virus across all lineages expanded (median 5.08× 10^8^ copies/ml +/− 2.74 × 10^8^; IQR 2.89 × 10^8^; Summarized in Supplementary Table I). To anchor these observations with a non-engineered cell line that closer mimics the *in vivo* setting, we subsequently tested all variants using Calu3 cells (Fig. 1D), which is a human lung adenocarcinoma-derived cell line. Titrations performed in Calu3 cells demonstrated viral titers in which pre-Omicrons and Omicrons clustered into high and low replication levels, respectively (Fig. 1D). Whilst a similar grouping profile of viral replication was observed in the VeroE6-T2 cell line, the latter could sustain levels of viral replication across all variants that were orders of magnitude higher, when compared to the Calu3 cell line (Fig. 1H). In an attempt to further resolve the entry requirements and the basis of susceptibility to infection across all lines, we stained for the SARS-CoV-2 entry factors ACE2 (Fig. 1I) and TMPRSS2 (Fig. 1J) and measured expression levels using flow cytometry. Whilst the HAT-24 line and VeroE6-T1sustained similar profiles of viral permissiveness, their receptor profiles could be readily contrasted as ACE2 high: TMPRSS2 low and ACE2 low: TMPRSS2 high respectively. Whilst the VeroE6-T2 line was defined as the line expressing the highest levels of TMPRSS2 but with low levels of ACE2, similar to VeroE6-T1 cells. The higher levels of TMPRSS2 expressed by VeroE6-T2 cells compared to the VeroE6-T1 cell line may provide the mechanistic basis of higher viral replication in the former line across all variants studied. Although simply higher levels of TMPRSS2 expression contradicts with that observed in the HAT-24 line with low TMPRSS2 expression and furthermore in observations of Omicron lineages attenuation in both cells and tissues that express high TMPRSS2 ^26^. Therefore, we cannot rule out the role other cellular factors and/or mechanisms beyond ACE2 and TMPRSS2 co-expression, that would enable high replication across both pre-Omicron and Omicron lineages in the VeroE6-T2 line.

**Figure 1.**
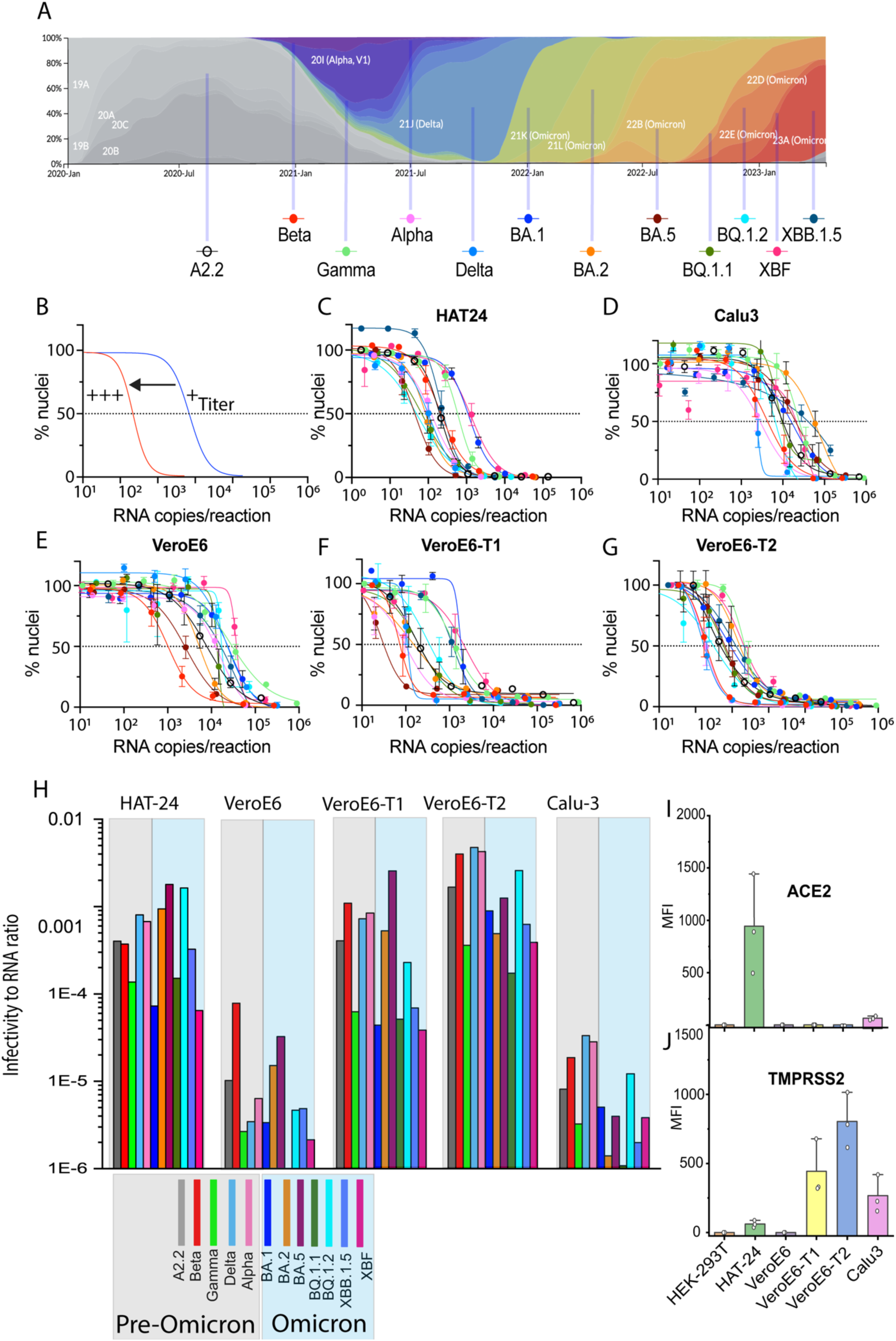
Engineered cell lines with TMPRSS2 reveal a continuum of sensitivity across all major primary SARS-CoV-2 variants. **A.** Globally dominant lineages presented over time and colored by Clade (nextstrain.org/sars-cov-2). The lineages listed below are the primary isolates used in this study which represent either variants of concern and/or dominant variants in the community at a specific time. **B.** Schematic illustrating high content datasets relative to RNA input. A shift in the sigmoidal curve to the left represents a viral isolate with a greater titer (low viral input can sustain significant cytopathic effects) which is readily enumerated by high content microscopy using live fluorescent nuclei counts. The dotted line represents 50% loss of nuclei and interpolated values can be used to determine viral titers. **C-G.** Titration curves per RNA input for the 12 primary clinical isolates highlighted in A. **C.** The Hek 239T engineered HAT-24 cell line co-expressing ACE2 and TMPRSS2, as previously described ^16^. **D.** Calu-3 human lung adenocarcinoma cell line. **E.** VeroE6 cell line. **F.** VeroE6 cell line expressing TMPRSS2 (VeroE6-T1), as previously described^18^. **G.** VeroE6 cell line expressing TMPRSS2 (VeroE6-T2), as previously described ^19^. Herein we refer to this line as VeroE6-T2. G. **H.** Infectivity is calculated by the viral dilution requited to sustain 50% of the loss of nuclei and then expressed as an infection to RNA ratio on the Y-axis (VE50). Grey and blue shading represents pre-Omicron and Omicron lineages, respectively. Infectivity to RNA ratios are derived from data generated from C-G. and is representative of three independent experiments. **I & J.** Cell surface staining and flow cytometry of cells used herein for **I.** TMPRSS2 and **J.** ACE2. Each data point represents an independent experiment.

### High ACE2 expression in the presence of high TMPRSS2 augments pre-Omicron lineages but attenuates Omicron lineages

To confirm that viral entry in the VeroE6-T2 cell line was primarily mediated by TMPRSS2-Spike S2 activation, virus was titrated in the presence of the TMPRSS2 inhibitor Nafamostat. Treatment decreased susceptibility to infection across all variants (Fig. 2A-D). In contrast, treatment of both VeroE6 and VeroE6-T2 cells with the Cathepsin L inhibitor, E64D, only inhibited pre-Omicron and Omicron viral entry in VeroE6 cells. (Fig. S1). Therefore, TMPRSS2 Spike S2 activation was still functional in VeroE6-T2 cells across pre-Omicron and Omicron lineages. We subsequently increased ACE2 expression in the VeroE6-T2 cell line (Fig. 2E) to determine if this would further augment viral replication across all variants. This led to augmented infection for all pre-Omicron lineages, especially Delta, which contrasted to the attenuated infection demonstrated across all Omicron lineages (Fig. 2F-I). Using this approach, Omicron lineages could be sub-grouped into lineages circulating in 2022, which showed less attenuation compared to that demonstrated with the more contemporary 2023 recombinants XBF and XBB.1.5 (Fig. 2I). Paradoxically, these two latter isolates have the highest level of ACE2 affinity as they both have the F486P Spike polymorphism, which has been shown to enable the greatest fitness in transmission ^28^. To confirm the efficiency of primarily ACE2 binding and entry for these specific lineages, we developed an ACE2 inducible cell line. In brief, this cell line is an important control at two levels. Firstly, this provides a setting where a viral receptor can be expressed at increasing levels, with the use of a drug-induced promoter like that outlined in the Affinofile assay developed by Lee and colleagues to resolve the entry requirements of HIV-1^29^. Secondly, in the absence of TMPRSS2, ACE2 becomes the dominant entry factor and activation of viral fusion is mediated by endosomal uptake and Cathepsin L cleavage of Spike. The latter control is important in uncoupling the complex activity of viral entry with respect to combined ACE2 and TMPRSS2 dynamics, through focusing entry requirements of each variant on primarily ACE2. Using the Affinofile system outlined herein, viral entry and replication of a primary isolate at low ACE2 expression would confirm a variant’s efficient use of ACE2 independent of TMPRSS2 activity. We focused on SARS-CoV-2 lineages (XBB.1, XBB.1.5, CH.1.1 and XBF) with known changes in Spike that have led to greater ACE2 affinity ^30^. The variants were chosen for three reasons. Firstly, they all represent Omicron lineages circulating at the same time. Secondly, the XBB.1.5 and XBF lineages have the F486P Spike polymorphism that is known to significantly increase ACE2 affinity. Finally, XBB.1 is the parent for XBB.1.5 with the only change in Spike F486P. Using this approach, we could readily resolve efficient ACE2 usage with lineages harboring the F486P Spike polymorphism, by comparing XBB.1 and CH.1.1 (486F) with XBB.1.5 and XBF (486P) (Fig. 2J & K). This result directly contrasts with the one observed in the VeroE6-T2-ACE2 line and therefore supports Omicron lineage-induced attenuation to be primarily related to the dynamics between ACE2 and TMPRSS2 in viral entry.

**Figure 2.**
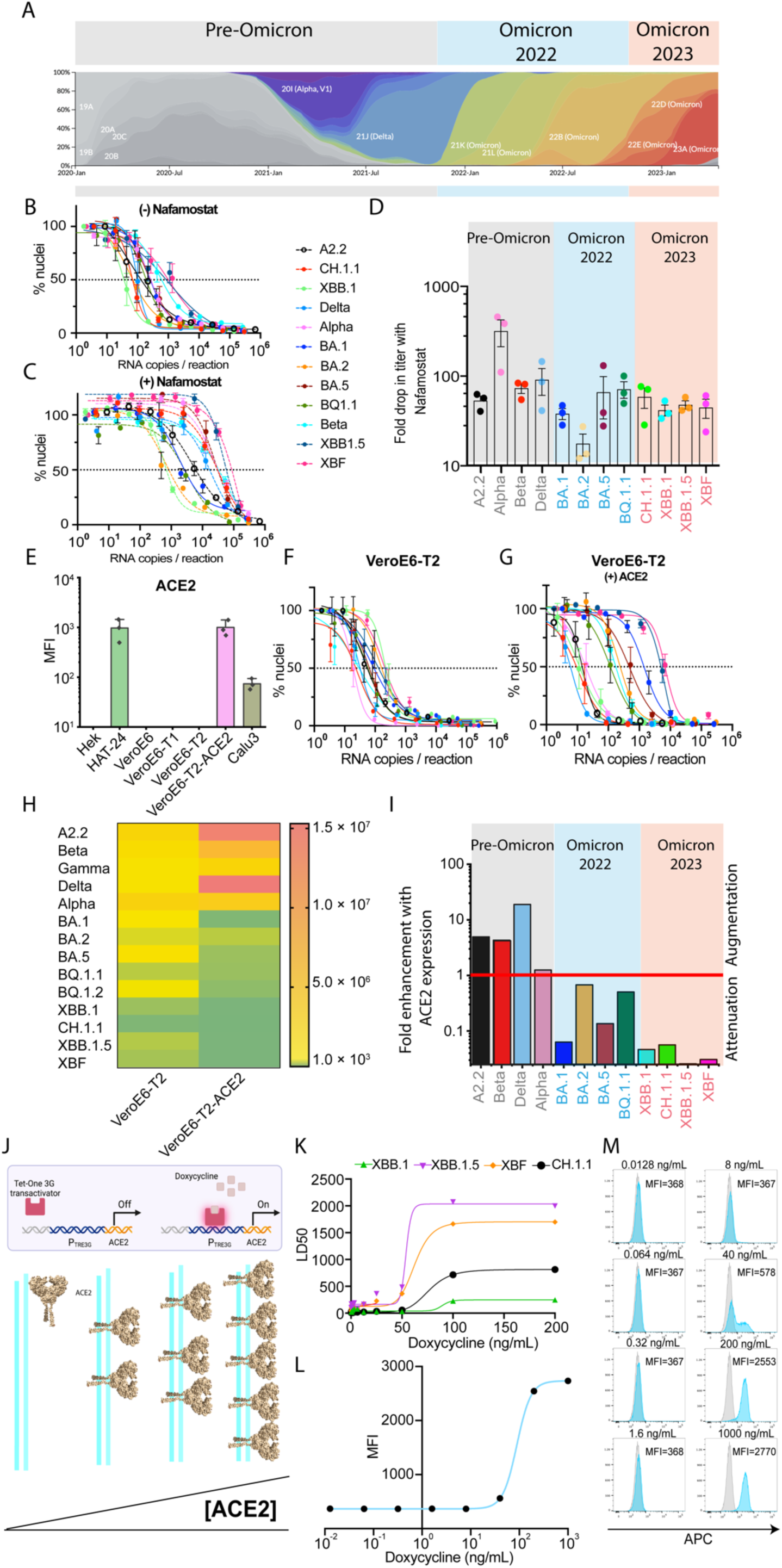
High co-expression of ACE2 and TMPRSS2 attenuates Omicron lineage use of TMPRSS2. **A.** Frequency of SARS-CoV-2 variants colored by lineage (nextstrain.org/sars-cov-2) over the pandemic. Three periods of viral spread are highlighted by shading in grey (Pre-Omicron), blue (2022 Omicron) and pink (2023 Omicron). **B-C.** Viral titrations of 12 primary SARS-CoV-2 isolates in the VeroE6-T2 cell line with B. vehicle control only (dimethyl sulfoxide; DMSO) or C. the TMPRSS2 inhibitor Nafamostat (20 µM) demonstrate a shift of the titration curves from the left to the right. This indicates reduced viral titers in the presence of Nafamostat, which provides a quantitative measure of TMPRSS2 dependence for entry in the VeroE6-T2 cell line. **D.** Summary of fold reduction in titers in the presence of Nafamostat from B-C; each data point represents an independent experiment. Each variant period is shaded in grey, blue and pink to align with variant periods outlined in A. All pre-Omicron lineages demonstrate high dependence on TMPRSS2, indicates by a significant reduction in titers in the presence of Nafamostat, there is a significant decrease in TMPRSS2 use observed with the arrival of Omicron BA.1 and BA.2. **E.** Surface expression of ACE2 in the VeroE6-T2-ACE2 cell line, measured using flow cytometry, relative to other cell lines presented in Figure 1. **F-G.** Viral titers of 12 primary SARS-CoV-2 isolates in the F. VeroE6-T2 cell line (low ACE2 expression) and G. VeroE6-T2-ACE2 cell line (high ACE2 expression). Left and right shifts in the sigmoidal curves indicate attenuation and augmentation of viral entry, respectively. **H.** Summary of viral titers from F-G. Note: pre-Omicron titers can be readily observed to be augmented in the high ACE2 and high TMPRSS2 setting. I. Ratios of VeroE6-T2-ACE2 to parental VeroE6-T2 titers establish levels of augmentation or attenuation across variant eras outlined in A. The red line is where titers are equivalent in each cell; augmentation is indicated by titer ratios above the red line whilst attenuation is represented by variants with titer ratios below the red line. **J.** Affinofile cell line comprised of HEK-293T cells engineered to express variable levels of ACE2 using a Doxycycline sensitive promoter. The greater the [Doxycycline], the higher the expression of ACE2. **K.** Viral titers for XBB.1, XBF, XBB.1.5 and CH.1.1 using the Affinofile cell line outlined in J. The F486P polymorphism in XBF and XBB.1.5 (known to increase ACE2 affinity) can be resolved in the Affinofile system with increased LD50 values (indicative of augmented viral entry and infection). **L-M**. Summary of ACE2 expression in the Affinofile system with increasing [Doxycycline], as measured using surface staining for ACE2 and flow cytometry. Blue histogram = Doxycycline-induced cells; grey histogram = non-induced cells. MFI = mean fluorescence intensity.

### Modification of the known ACE2-TMPRSS2 binding domain at the ACE2 CLD rescues TMPRSS2 use across all SARS-CoV-2 Omicron lineages

The complexity of SARS-CoV-2 entry requirements could relate to the distinct and often tissue-specific roles of ACE2 *in vivo*. ACE2 is physiologically important in two distinct pathways. The first relates to the carboxypeptidase activity of ACE2 and the need to cycle from membrane bound to soluble forms to regulate RAAS ^31^. In this setting, a disintegrin and metalloprotease 17 (ADAM17) and TMPRSS2 regulate the shedding and retention, respectively, through differential ACE2 cleavage in the exposed neck of ACE2. ACE2 has a second physiological role as a molecular chaperone, involving the heterodimerization of ACE2 with one of two solute carrier proteins, SLC6A19 and SLC6A20. It has been previously demonstrated that SARS-CoV-1 infectivity can be augmented by the presence of TMPRSS2 -ACE2 complexes facilitated by the ACE2 CLD domain as part of RAAS regulation ^17^. This led us to investigate if SARS-CoV-2 entry requirements are also regulated by the ACE2 CLD and its ability to recruit partners like TMPRSS2. Given the role of the ACE CLD and TMPRSS2 in RAAS regulation is through blocking of ACE2 shedding, we pragmatically investigated this process through initially screening several different cell lines for soluble ACE2 enzymatic activity (a surrogate for ACE2 shedding) in cell line supernatants ^30^. Detectable ACE2 activity was observed across all engineered cell lines except the VeroE6-T2-ACE2 cell line (Fig. 3A-B). This observation is consistent with a previous study that highlights TMPRSS2 competing with other proteases to cleave ACE2 in a manner that impedes subsequent release of soluble ACE2^17^. This was in line with our observations in the VeroE6-T2-ACE2 line and furthermore supports a phenotype where no ACE2 is available for shedding in this line. However, the high level of ACE2 expression alongside TMPRSS2 on the VeroE6 cell surface, is not consistent with that observed in many studies using HEK-293T cell lines ^17,32^. Rather, the latter cell line promotes extensive ACE2 cleavage by TMPRSS2 and significantly reduces the levels of ACE2 from the cell surface. Throughout, we have C-terminally tagged ACE2 with the C9 tag, with the objective to observe ACE2 cleavage when co-expressed with TMPRSS2. In stark contrast to that observed in previous studies with prior observations within the HEK293T cell line ^17,32^, we observed high ACE2 co-expression with TMPRSS2 in the VeroE6 cell line to sustain primarily intact and uncleaved ACE2 (Fig. 3D). Therefore, the unique entry phenotypes observed using the VeroE6-T2-ACE2 cell line are consistent with high levels of ACE2 matching TMPRSS2 at the surface.

**Figure 3.**
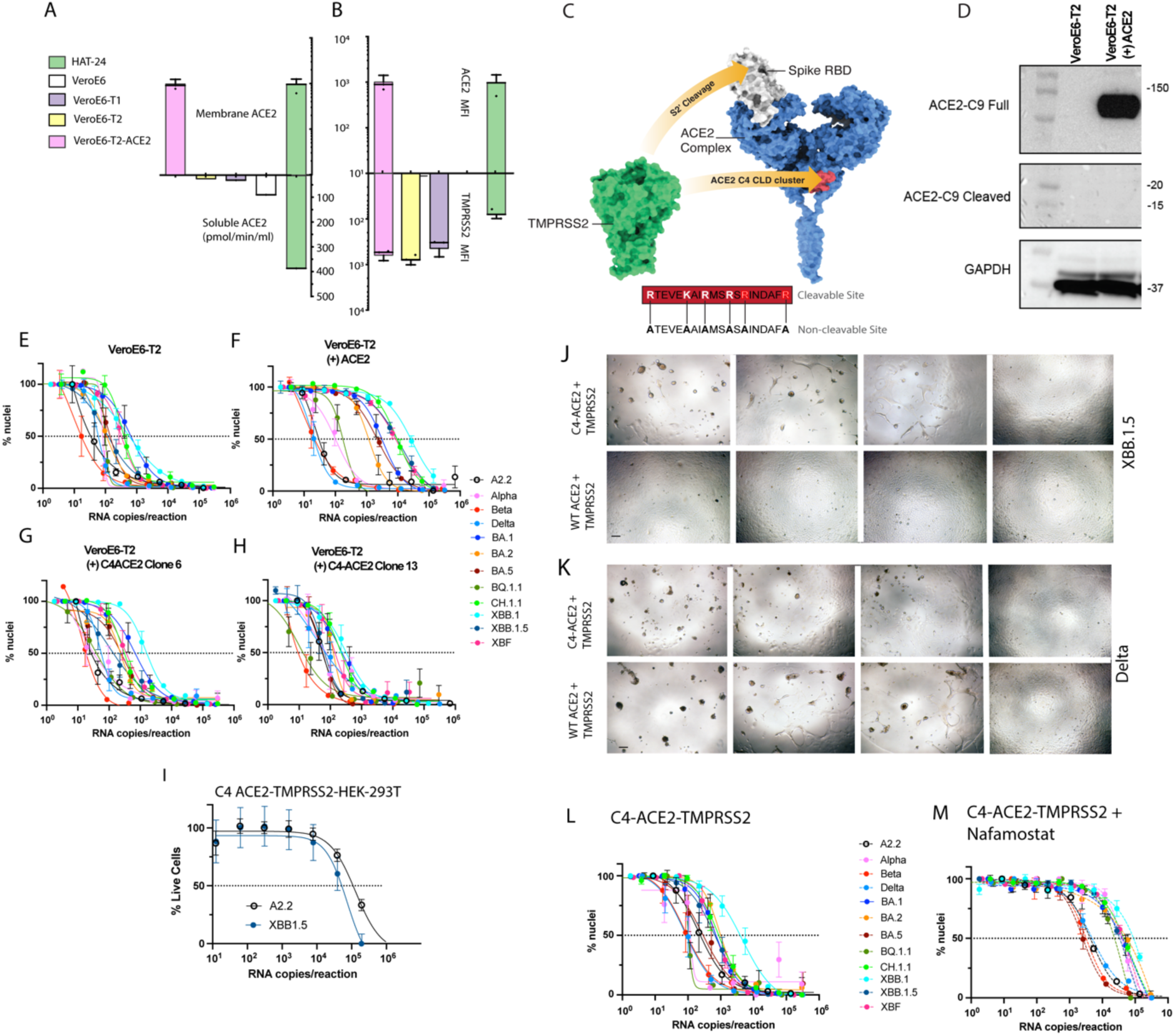
TMPRSS2 cleavage of ACE2 resolves the differential entry requirements for pre-Omicron and Omicron lineages. **A.** Membrane and soluble ACE2 levels in engineered cell lines. Membrane ACE2 is detached through flow cytometry and represented as mean fluorescence intensity. The activity of soluble ACE2 is detected using the supernatant of 80% confluent cell lines after three days of culture, as previously described^1^. **B.** Ratios of cell surface ACE2 and TMPRSS2 detected via flow cytometry as per A. **C.** Left: Schematic of Spike (grey), TMPRSS2 (blue) and ACE2 (yellow). Right: ACE2 structure with known TMPRSS2 binding site in red (Structure PDB 6M17) and the sequence of the C4-ACE2 mutant to the right in black with alanine substitutions in bold. **E-H.** Viral titration curves across 12 primary SARS-CoV-2 isolates with **E.** low WT ACE2: High TMPRSS2, **F.** high WT ACE2:High TMPRSS2, **G.** NC-ACE2:High TMPRSS2 (clone 6 with lower NC-ACE2 expression to clone 13 in G.) and **H.** high NC-ACE2:High TMPRSS2. Input viral inoculate in E-H is presented here in RNA copies per infection/reaction**. I-J.** Contrasting replication across variants appearing in WT ACE2:High TMPRSS2 and NC-ACE2:High TMPRSS2 cell lines presented in F. and H., respectively, for pre-Omicron lineage I. Delta, and the Omicron lineage J. XBB.1.5. Fields of view from left to right are starting at 1/2000 dilution with 1/5 dilution steps in each successive frame. The Omicron lineage XBB.1.5 leads to extensive viral syncytia formation only in the presence of NC-ACE2, whilst the pre-Omicron Delta demonstrates viral syncytia at a similar level across WT and NC-ACE2 engineered cell lines. Scale bar is at 20 µM. K-L. Using the cell clone presented in H. we titrated virus in the **K.** absence or **L.** presence of saturating levels of the TMPRSS2 inhibitor Nafamostat. Data presented in K-L has been input as RNA copies of virus per variant. **M.** Replication of pre-Omicron and and Omicron lineages using C4-ACE2 and TMPRSS2 in a Hek-293T engineered cell line. **N.** Increasing expression of WT ACE2 with constant TMPRSS2 only facilitates entry with respect to the pre-Omicron clade A2.2.

With the majority of studies on SARS-CoV-1 and SARS-CoV-2 entry using the HEK-293T cell line, many molecular events have been attributed to TMPRSS2’s ability to cleave ACE2 and in doing so augmenting viral entry and fusion. Whilst we also observe augmentation of entry in the VeroE6-T2-ACE2 cell line using early SARS-CoV-2 clades, this is in the context of intact uncleaved ACE2 expressed at the surface at equivalent high levels to TMPRSS2. So the alternate hypothesis here may not be ACE2 cleavage, but rather the ability of ACE2 and TMPRSS2 to form a complex that subsequently influences Spike S2 cleavage. Depending on the cell type this may also coincide with (in the HEK293T Line) or without (in the VeroE6 line) ACE2 cleavage. To uncouple ACE2 from TMPRSS2 we generated the C4-ACE2 mutant known to be resistant to TMPRSS2 cleavage when expressed in the HEK-293T cell line (Fig. 3C)^17^. Curiously C4-ACE2 mutant removes a cluster of positive amino acid residues (Arg and Lys) upstream of the known ACE2 cleavage site^17^. So rather than representing an ACE2 that has the TMPRSS2 removed, the C4-ACE2 mutant may simply represent a mutant of ACE2 that is uncoupled from TMPRSS2 both physically and enzymatically. We expressed the C4-ACE2 mutant to equivalent levels to (WT) ACE2 that is observed in the VeroE6-T2-ACE2 cell line (Fig. S2). Viral titrations using the C4-ACE2 mutant expressed alongside high levels of TMPRSS2 in a VeroE6 cell line background demonstrated lower levels of augmentation of pre-Omicron variants in the C4-ACE2 cell line compared to WT ACE2, with the largest drop observed in the earliest circulating Clade A.2.2 lineage. Given WT ACE2 is uncleaved in the VeroE6-T2-ACE2 line this is consistent with a mechanism of SARS-CoV-2-augmented entry being related to a ACE2-TMPRSS2 complex rather than TMPRSS2 cleavage of ACE2^17^. In contrast, Omicron lineages were able to infect C4-ACE2 cells effectively, and as such, all major SARS-CoV-2 lineages to date were able to maintain viral replication at similar levels (Fig. 3G, 3H). To determine if the C4-ACE2 phenotype could be reproduced in another cell line, we generated a HEK-239T line that only expresses NC-ACE2 and TMPRSS2, but not WT ACE2. To consolidate testing, we compared the pre-Omicron lineage A.2.2 versus the contemporary Omicron lineage XBB.1.5. To reproduce the C4-ACE2 phenotype, equivalent permissiveness to both clades (one that emerged in 2020 and the other in 2023) should proceed and this was observed (Fig. 3I). Therefore, these results suggest that C4-ACE2 sustained replication of all SARS-CoV-2 variants tested at equivalent levels, whereas WT ACE2 (with the presence of high TMPRSS2) only provides an entry advantage for early pre-Omicron lineages.

To resolve the mechanism of replication of omicron variants we compared the early pre-Omicron A.2.2 and the more contemporary XBB.1.5 omicron over time. Within 2 days of culture XBB.1.5 Omicron cultures accumulated significant syncytia only in the VeroE6-T2 C4-ACE2 setting (3 J.). This was similar to that observed in Delta (Fig. 3K.), with the exception Delta could generate syncytia across both WT ACE2 and C4-ACE2 engineered VeroE6-T2 lines. The extensive syncytia observed only in XBB.1.5 cultures only with C4-ACE2 pools and TMPRSS2 is consistent with TMPRSS2-Spike S2 cleavage and activation of viral fusion in this setting. Thus, uncoupling TMPRSS2 from ACE2 through the introduction of the C4 mutation in ACE2 then may sustain TMPRSS2 in a state that is able to engage Omicron Spike S2 cleavage. To test if infectivity across all variants is TMPRSS2 dependent with C4-ACE2 pools, we titrated virus in the presence and absence of the TMPRSS2 inhibitor Nafamostat using the C4-ACE2 VeroE6-T2 cell line. Following treatment, large inhibitory shifts in viral titer curves were observed, with all Omicron lineages shifting approximately 100-fold lower in titer when using the C4-ACE2 VeroE6-T2 cell line (Fig. 3L-M). Thus, we conclude that TMPRSS2 association with ACE2 at the CLD may act as an “off” switch for subsequent TMPRSS2 activation of Omicron Spikes whilst pools of ACE2 separated from TMPRSS2 through modification of the ACE2 C4 cluster within the CLD sustain an “on” switch configuration (summarized in graphical abstract). Whilst early lineages of SARS-CoV-2 favour ACE2-TMPRSS2 complexes (as does SARS-CoV-1), it is clear that all Omicron lineages no longer align with this entry pathway.

### SARS-CoV-2 lineages over the last 4 years of the pandemic have evolved towards on entry pathway of only C4-ACE2 and TMPRSS2 use

As observed above, early pre-omicron lineages consistently replicated across both WT and C4 ACE2 pools and did so with efficient TMPRSS2 usage. In contrast Omicron lineages were observed primarily to utilize C4-ACE2 and TMPRSS2. In an entry requirement setting, pre-Omicrons have molecular tropism consistent with dual/promiscuous tropism, whilst Omicron lineages have a molecular tropism profile that is consolidated towards the C4-ACE2 & TMPRSS2 entry phenotype. To resolve this further, we determined the augmentation/attenuation profiles of every major variant over the last 4 years. This covered all major pre-Omicron lineages and every major Omicron lineage starting from 2022 and concluding in 2024. Using this approach we observed groupings of SARS-CoV-2 lineages based on augmentation or attenuation when WT ACE2 and TMPRSS2 was expressed at equivalent levels within the VeroE6 cell line (VeroE6-T2-ACE2). Whilst pre-Omicrons were consistently augmented (with the exception of Alpha), each year of Omicron variants were observed to be consistently attenuated in their ability to enter the latter cell line. Presently, both JN.1 and the sub-lineage KP.3 represent the peak of this attenuation (4.A & B). In contrast, the use of C4-ACE2 and TMPRSS2 is sustained across all variants. Whilst there is an upward trend in augmentation over time for combined C4 ACE2 and TMPRSS2 use, only KP.3 is presently significantly greater than the earliest pre-Omicron clade A2.2 in this setting (4.6 fold versus 1.38 augmention in the presence of C4 ACE2 compared to the parental VeroE6-T2 line for KP.3 and Ancestral clade A.2.2 respectively; *p* = 0.0048).

**Figure 4.**
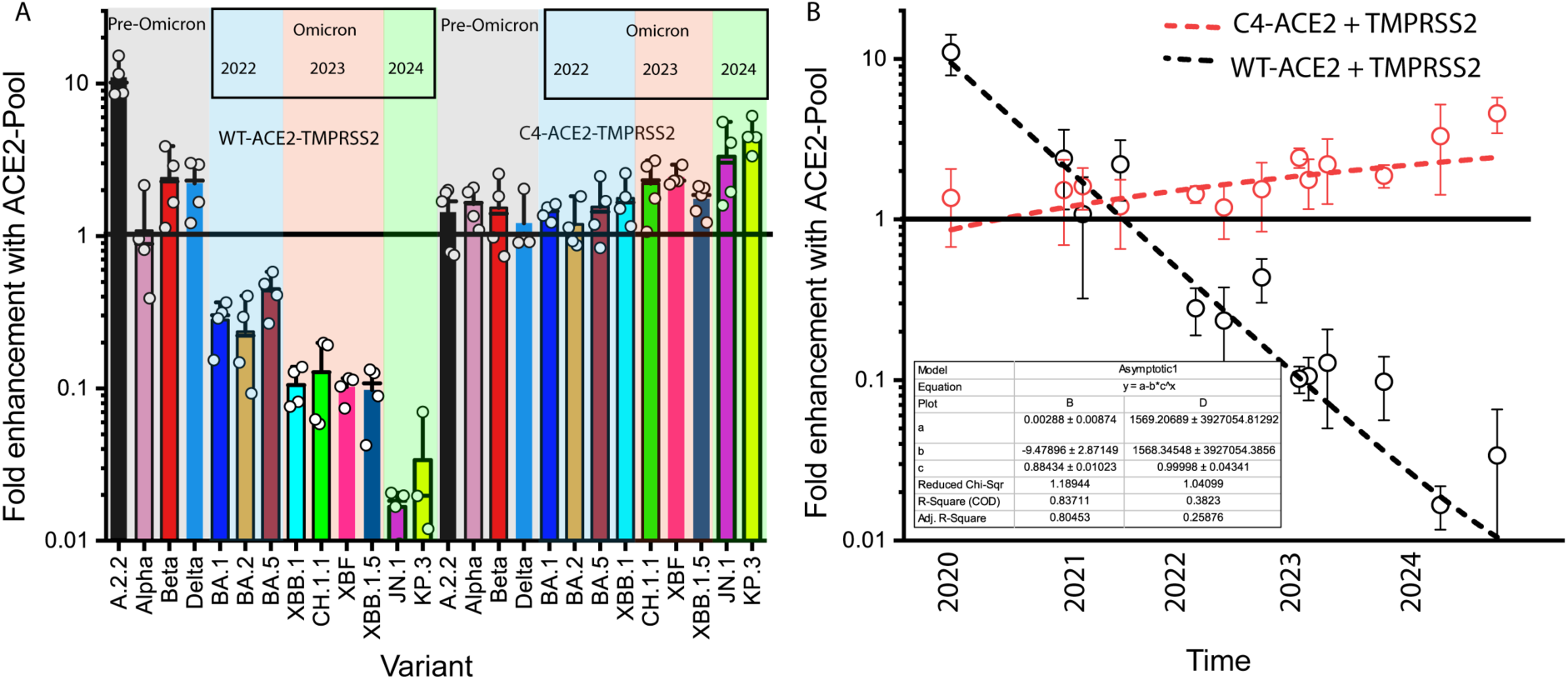
**A&B.** Summary of fold augmentation and attenuation of all major variants to date across the two pools of ACE2 (WT ACE and C4-ACE2 when co-expressed with TMPRSS2 in a VeroE6 background. Fold attenuation or augmentation is calculated as the fold change of the engineered cell line expressing each ACE2 pool (C4-ACE2 VeroE6-T2-See panel Fig.3 H.) compared to the parental expressing TMPRSS2 and low ACE2 (VeroE6-T2). In A. data points are presented from independent experiments. B. Summarizes this over time with respect to the pandemic. Mean and Standard deviations are calculated from the 4 independent experiments presented in A.

### ACE2 oligomerization with Solute Carriers SLC6A19 and SLC6A20 occupies the ACE2 CLD and may represent the physiological pool of ACE2 that can uncouple the ACE2-TMPRSS2 complex in a manner similar to C4-ACE2

The continued use of TMPRSS2 by Omicron lineages could explain the discrepancies in Omicron replication observed *in vivo* versus that observed *in vitro* by ourselves and others ^3,16,26,33^. For instance, overexpression of ACE2 and TMPRSS2 in many cell lines readily divides pre-Omicron and Omicron lineages by biasing of cells towards ACE2 pools complexed with TMPRSS2. Whilst the latter may reflect the attenuation in the lung, other tissue sites *in vivo* can readily sustain efficient SARS CoV-2 replication ^4,34,35^. The key to further elucidating Omicron tropism *in vivo* is to understand the two very separate roles ACE2 plays *in vivo,* and how this potentially relates to Omicron tropism. Whilst the abovementioned the role of ACE2 in the RAAS pathway and in the lung are well established, the majority of ACE2 within the body works as a chaperone of several solute carrier (SLC) proteins. This physiological role for ACE2 aligns to our study for three reasons. Firstly, SLCs and ACE2 heterodimerize which results in structural occupancy of the TMPRSS2 binding site at the ACE2 dimer interface (ACE2 amino acid residues 710 and residue 716). Secondly, ACE2 in its role as a chaperone would need to remain membrane bound and would not benefit from the proteolytic regulation by TMPRSS2 and/or ADAM-17 observed in the RAAS pathway. Finally, modelling studies of ACE2 dimers complexed with SLC6A19 have revealed that TMPRSS2 is obstructed from accessing the C4 CLD site^36^ and thus ACE2 complexed to solute carriers like SLC6A19 would keep TMPRSS2 structurally at a distance in a manner the C4-ACE2 mutant would mimic.

Given the structure of ACE2 in its role as a chaperone for SLCs has the potential to rescue Omicron infection, we then investigated the expression of SLC6A19 and SLC6A20 across various tissue targets of SARS-CoV-2 throughout the body to resolve their potential in driving SARS-CoV-2 tropism *in vivo*. We analyzed single cell RNAseq data from the nasal cavity, lung and ileum, as they cover all major SARS-CoV-2 reservoirs *in vivo*. Here we compared the ratios of ACE2 and TMPRSS2, alongside ACE2 partners SLC6A19 and SLC6A20. Ciliated nasal epithelial cells (HNCs) expressed low levels of ACE2 and were associated with low SLC6A20 and moderate TMPRSS2 expression (Fig. 5A, B and G). In the lung, TMPRSS2 dominated expression over ACE2 and SLC6A20 in type I & II pneumocytes (Fig. 5C, D and H). In contrast, in enterocytes of the ileum, ACE2 and SLC6A19 were expressed at equivalent high levels, whilst TMPRSS2 levels were lower than that observed in lung pneumocytes (Fig. 5E, F and I). SLC6A20 was also present at low levels within known SARS-CoV-2 cellular targets. Therefore, *in vivo,* it is apparent that the lung and the lower intestine represented the two differing compartments of ACE2: i.e. RAAS and regulation of soluble ACE2 in the lung and SLC6A19 chaperone function in the intestine. The latter is consistent with *in vivo* observations associated of ACE2 expression in the intestine being essential in regulating SLC6A19 surface expression ^37,38^.

**Figure 5.**
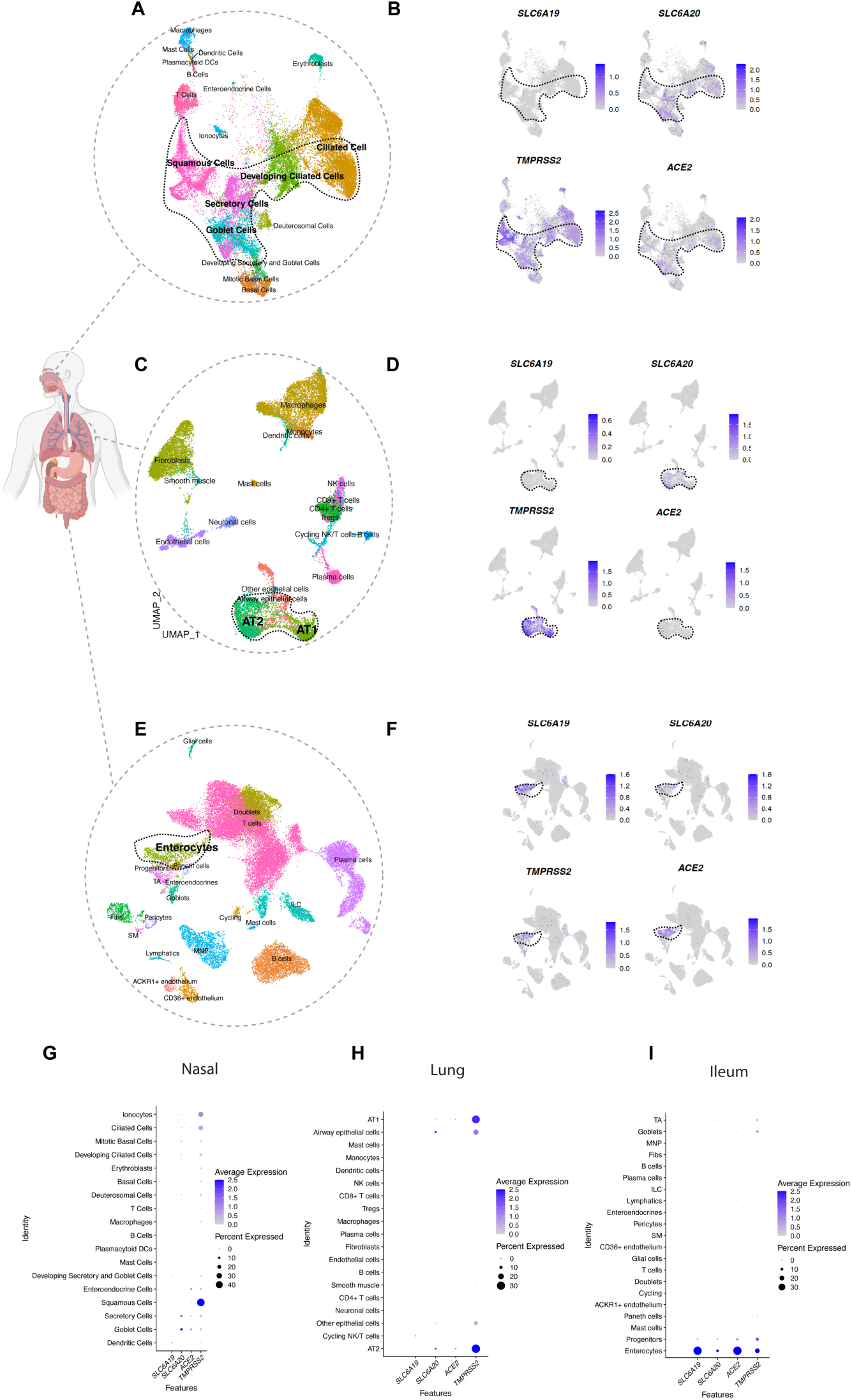
Expression of known SARS CoV-2 viral receptors with solute protein carriers SLC6A19 and SLC6A20 in targets of the nasal epithelium, lung and lower intestine. **A.-F.** Uniform manifold approximation and projection (UMAP) of **A.** Nasal Epithelia. **C.** Lung and **E.** Small intestine using single nuclei RNA sequencing. Cell types are color-coded. Target cells within each tissue are highlighted in B. D. & F. in respective tissues, with expression of protein solute carriers SLC6A19 and SLC6A20 alongside the known SARS CoV-2 receptors ACE2 and TMPRSS2. In **G.** to **I.** Relative expression of each solute carrier with ACE2 & TMPRSS2 is presented for each major cell type resolved in each tissue.

## Discussion

Resolution of viral entry pathways is essential to track the fitness of viral variants and their ability to spread, and to understand the pathogenesis and potential clinical manifestations as they appear in the community. In most instances the shifts in viral entry are subtle, however there are occasions where a significant accumulation of changes in a viral glycoprotein can enable a seismic shift in how the virus engages and infects cells and tissues. Whilst most SARS-CoV-2 infections are rapidly cleared, prolonged viral replication in immunocompromised or immunosuppressed patients has been shown to lead to an accumulation of amino acid substitutions or deletions within Spike ^39–42^. This accumulation of mutations is the primary hypothesis for the emergence of new immune-escape variants, such as the VOC Omicron. ^41^. As with other chronic viral infections, SARS-CoV-2 has evolved to target cells and tissues in a manner different from earlier circulating lineages. From preliminary studies by ourselves and others, it was immediately apparent that Omicron lineages were engaging the serine protease TMPRSS2 differently than Delta and earlier variants ^16,26,33^. This change in entry manifested *in vivo* in animal models ^3^, as well as in a marked decrease in COVID-19-associated pneumonia prevalence in patients infected with Omicron. ^43^. Herein, using extensive panels of genetically intact primary isolates from early 2020 through to early 2024, we have mechanistically dissected the evolving entry requirements of SARS-CoV-2 variants across the pandemic. Using this approach, we observed that the major shift in entry requirements from Delta to Omicron lineages is the culmination of TMPRSS2 activity on both ACE2 and SARS-CoV-2 Spike. If the known TMPRSS2 C4 CLD interaction site of ACE2 is available, then latter activation of Omicron Spike by TMPRSS2 does not proceed (ACE2 renders TMPRSS2 in the “off” conformation and as such does not enable Omicron Spike activation through S2 cleavage). In contrast, pre-Omicron lineages are augmented under the same entry conditions.

The phenotypic shift from Delta to Omicron has been studied extensively and the changing requirements for entry have been documented. The Omicron entry hypothesis is broadly based on the inefficient cleavage of the S1/S2 Spike domains, resulting in increased non-cleaved S1/S2, and reduced fusogenic activation of S2 by TMPRSS2 and increased endosomal entry pathways. In the endosomal entry pathway, Spike utilizes the cysteine protease Cathepsin L to activate and drive S2 fusion across the endosomal membrane ^26,33^ which does not require prior S1/S2 Spike cleavage by Furin. Whilst replication of Omicron BA.1 supports this hypothesis, post-BA.1, there has been an emergence of subsequent lineages with a continuum of S1/S2 cleavage ^44,45^. To date, this has not correlated with an entry phenotype consistent with that observed for pre-Omicron lineages such as Delta. Our study confirms that Omicron lineages may indeed be forced into using the endosomal pathway when TMPRSS2 Spike activation is attenuated or switched off. Whilst TMPRSS2-independent entry through endocytosis is a well-documented pathway *in vitro*, accumulating evidence *in vivo* supports continued use of TMPRSS2 as a dominant entry pathway required for viral replication and persistence of Omicron lineages *in vivo* ^2,34^. In addition, given the fitness gain of the virus that uses TMPRSS2, ongoing use of this serine protease for efficient viral entry is consistent with its high transmissibility *in vivo*.

Rather than a shift to a primary endocytosis entry pathway, our results support the major shift of Omicron entry requirements at the plasma membrane to be conditions which uncouple TMPRSS2 from ACE2. To reconcile to disparate observations of Omicron entry requirements *in vitro* and *in vivo*, we propose a molecular tropism model based on “ACE2 pools” to phenotypically resolve the entry differences between Omicron lineages versus pre-Omicron lineages. In this model, under conditions that drive TMPRSS2-ACE2 association at the ACE2 CLD, there is augmentation of pre-Omicron lineages but progressive attenuation of Omicron lineages spanning from BA.1 to the more contemporary variants JN.1 and KP.3. The physiological relevance of the proposed model is consistent with the pools of ACE2 needed for the RAAS pathway where the CLD is available for TMPRSS2 and ADAM17 binding events versus pools of ACE2 that either play a chaperone role for many solute carrier proteins or are sustained in a conformation that do not enable TMPRSS2 binding (ACE2 pools excluded from TMPRSS2 binding). Evidence supporting ACE2’s chaperone role excluding TMPRSS2 -ACE2 CLD binding is observed not only in molecular docking studies of ACE2 complexed to the solute carrier protein SLC6A19, but across studies that have solved ACE2-SLC6A19 and ACE2-SLC6A20 structures (refs). Collectively these studies observe a lack of TMPRSS2 access through occupation of the ACE2 CLD C4 cluster through the formation of a dimer of heterodimers between ACE2 and its respective solute carriers (SLC6A19 or SLC6A20). Whilst SLC6A19 expression is enriched in the ileum and represents the ACE2 partner in this tissue, in other sites this protein is absent. An alternative SLC that is known to oligomerize with ACE2 is SLC6A20, which is expressed alongside ACE2 across all known SARS-CoV-2 primary tissue targets, ranging from the upper respiratory tract, lung, small intestine and brain (choroid plexus) ^46–48^. The above highlights a continuum of ACE2 chaperone roles across many tissues associated with SARS-CoV-2 infection ^49–53^. Whilst viral tropism across tissue sites such as the lung and the small intestine can be readily reconciled using the above ACE2 pool model, infection across the entire respiratory tract appears of greater complexity. Whilst high levels of TMPRSS2 in the lower respiratory tract increases the potential of ACE2-TMPRSS2 complexes and is consistent with omicron attenuation in the lung, the counter explanation of augmented replication within the nasal epithelium requires future investigation. Lower TMPRSS2 to ACE2 ratios, the availability for ACE2 to be engaged by TMPRSS2 at its CLD and the potential role of SLC6A20 for the latter, all need consideration. As Omicron infection proceeds *in vivo* in these compartments in a TMPRSS2-dependent manner^2^, we would support mechanisms that promote ACE2 pools to persist uncoupled from TMPRSS2 here.

As the high level of spread of SARS-CoV-2 globally continues and is compounded by the risk of co-circulation in other hosts, phenotypic surveillance of variants is needed to determine the consequences of continuing viral evolution. Rapid detection of shifts in viral tissue tropism and linkage to severe clinical presentation are important in real-tracking many viruses including SARS-CoV-2. However, to enable detection of tropism shifts, mechanistic resolution of the changing entry requirements is needed to distill this knowledge into entry screening platforms. Herein we have determined the key change in entry requirements, and also outlined assays that can readily determine how an emerging variant benefits from distinct ACE2 pools and importantly if it is on a trajectory towards or away from viral entry requirements that are now well known for SARS-CoV-1 and early circulating clades of SARS-CoV-2. Rapid phenotypic surveillance of potential variants or ACE2 using coronaviruses, may be key in identifying those which have higher clinical severity scores or tissue-specific clinical manifestations such as lung-related pneumonia. Furthermore, our observations provide the mechanistic foundation that can lead to alternate animal models, such as transgenic mice with ACE2 uncoupled from TMPRSS2, to better study Omicron lineages. This work further highlights the need to understand ACE2 pools at greater resolution and determine how the role of ACE2 as a solute carrier protein chaperone may be driving the evolution of SARS-CoV-2 entry requirements. At a minimum, this work enables platforms *in vitro* that can increase sensitivity of SARS-CoV-2 isolation, detection and phenotypic monitoring for pre-Omicron, current Omicron lineages, and importantly emerging Omicron variants. This will have a significant impact in diagnostics and future surveillance and management of this pandemic virus.

## Methods

### Cell culture

HEK293T cells (thermo Fisher, R70007), HEK293T derivatives including HAT-24 ^16^, VeroE6-T1 (CellBank Australia, JCRB1819), VeroE6-T2 ^19^, VeroE6-T2-ACE2 and VeroE6-T2-NC-ACE2 were cultured in Dulbecco’s Modified Eagle Medium (DMEM; Gibco, 11995073) with 10% fetal bovine serum (FBS) (Sigma; F7524). HEK293T-TetOne-ACE2 cells were cultured in DMEM with 10% fetal bovine serum (FBS) with 0.05µg/mL puromycin (Sigma; P8833). VeroE6 cells (ATCC CRL-1586) were cultured in Minimum Essential Medium (Gibco) with 10% FBS, 5% penicillin-streptomycin (Gibco; 15140122) and 1% L-glutamine (insert ref number). Calu3 cells were maintained in DMEM/F12 media (Gibco;12634010) supplemented with 1% MEM non essential amino acids (Sigma, M7145), 1% Glutamax (Gibco; 35050061) and 10% FBS. All cells used herein were only cultured and used within a range of 20 passages.

### Generation of stable cell lines

For generating VeroE6-T2 cells that stably express non-cleavable ACE2, expression plasmid pRRLsinPPT.CMV.GFP.WPRE was first modified to carry a multiple cloning site (MCS) at the 3’ end of GFP open reading frame (ORF) using Age1 and Sal1 cut sites. The non-cleavable form of hACE2 (NC-ACE2) carrying mutations described previously by Heurich et al ^17^was synthesized as a synthetic gBlock (IDT) and shuttled into the above plasmid using Xba1/Xho1 cut sites thus replacing the GFP ORF with NC-ACE2.

For generating HEK293T-TetOne-ACE2 cells, expression plasmid pLVX Tet-One Puro (Clontech) was modified to have a MCS at the 3’end of Tet responsive promoter TRE3GS using EcoR1/BamH1 cut sites. ACE2 ORF was amplified from pRRLsinPPT.CMV.ACE2.WPRE plasmid ^16^ and shuttled into pLVX Tet-One Puro-MCS expression plasmid using Not1/Xho1 sites to generate into pLVX Tet-One Puro-ACE2 plasmid. Plasmid sequences were validated by Nanopore Sequencing with the Rapid Barcoding Kit 96 (Oxford Nanopore Technologies) using the manufacturer’s protocol. The sequencing data was exported as FASTQ files and analyzed using Geneious Prime (v22.2). Open reading frames were then verified using Sanger Sequencing.

Cells expressing NC-ACE2 and inducible ACE2 were generated by lentiviral transductions as previously described ^16^. Briefly, lentiviral particles expressing the above genes were produced by co-transfecting expression plasmids individually with a 2nd generation lentiviral packaging construct psPAX2 (courtesy of Dr Didier Trono through NIH AIDS repository) and VSVG plasmid pMD2.G (Addgene, 2259) in HEK293T producer cells using polyethyleneimine as previously described ^54^.Virus supernatant was collected 72 h post-transfection, pre-cleared of cellular debris and centrifuged at 28,000 × g for 90 min at 4°C to generate concentrated virus stocks. Lentiviral transductions were then performed on VeroE6-T2 cells to generate VeroE6-T2-NC-ACE2 cells and HEK293T cells to generate HEK293T-TetOne-ACE2. The expression of ACE2 in these cells was induced with 200 ng/ml of Doxycycline (Sigma, D9891) as per the manufacturer’s instructions (ClonTech). Screening of clones was based on expression of high levels of ACE2 following Doxycycline in addition to being highly susceptible to infection with the early clade SARS-CoV-2 isolate A.2.2. HEK293T-TetOne-ACE2 cells were further modified to express hTMPRSS2 ^19^ by lentiviral transductions to generate HEK293T-TetOne-ACE2-TMPRSS2 cells.

## ACE2 Affinofile Assay

Cells, engineered with Doxycycline induced ACE2, were trypsinised and seeded as suspension at 5 × 10^3^ cells/well in 384-well plates. Cells were exposed to varying amounts of Doxycycline for a minimum of four hours and at then inoculated with SARS-CoV2 variants for 72 hours.

Following this, cells were stained live with 5% v/v nuclear dye (Invitrogen, R37605) and whole-well nuclei counts were determined with an INCell Analyzer 2500HS high-content microscope and IN Carta analysis software (Cytiva, USA). Data was normalized to generate sigmoidal dose–response curves (average counts for mock-infected controls = 100%, and average counts for highest viral concentration = 0%) and median Virus Effective (VE_50_) value for each doxycycline concentration was obtained with GraphPad Prism software ^25^

### Viral isolation, propagation and titration from primary specimens

SARS-CoV-2 variants were isolated from diagnostic respiratory specimens as previously described ^25^. Briefly, specimens testing positive for SARS-CoV-2 (RT-qPCR, Seegene Allplex SARS-CoV-2) were sterile-filtered through 0.22 µm column-filters at 10,000 x *g* and serially diluted (1:3) on HAT-24 cells (5 × 10^3^ cells/well in 384-well plates). Upon confirmation of cytopathic effect by light microscopy, 300 μL pooled culture supernatant and trypsinised cells from infected wells (passage 1) were added initially to a pellet of VeroE6-T2 cells (0.5 × 10^6^ cells) for 30 minutes and then subsequently transferred to a 6-well plate (well with 2 mL of MEM-2%FBS final) and incubated for 48 h to 72 h (or until cytopathic effects had led to loss of >50% of the cell monolayer). The supernatant was cleared by centrifugation (2000 x *g* for 10 minutes), frozen at −80°C (passage 2), then thawed and titrated to determine median 50% Tissue Culture Infectious Dose (TCID_50_/mL) on VeroE6-T2 cells according to the Spearman-Karber method ^55^. Viral stocks used in this study correspond to passage 3 virus, which were generated by infecting VeroE6-T2 cells at MOI=0.025 and incubating for 24 h before collecting, clearing, and freezing the supernatant as above in 100 µl aliquots. Sequence identity and integrity were confirmed for both passage 1 and passage 3 virus via whole-genome viral sequencing using an amplicon-based Illumina sequencing approach, as previously described ^56^. The latter was also used in parallel for sequencing of primary nasopharyngeal swabs.

Virus titrations were carried out by serially diluting virus stocks (1:5) in MEM-2%FBS, mixing with cells initially in suspension at 5 × 10^3^ cells/well in 384-well plates and then further incubating for 72 h. The cells were then stained live with 5% v/v nuclear dye (Invitrogen, R37605) and whole-well nuclei counts were determined with an IN Cell Analyzer 2500HS high-content microscope and IN Carta analysis software (Cytiva, USA). Data was normalized to generate sigmoidal dose–response curves (average counts for mock-infected controls = 100%, and average counts for highest viral concentration = 0%) and median Virus Effective (VE_50_) values were obtained with GraphPad Prism software ^25^. To assess the TMPRSS2 usage of the virus isolates, titrations on VeroE6-T2 were performed in the presence of saturating amounts of Nafamostat (20 μM). Titrations performed in parallel in equivalent volumes of DMSO served as controls and were used to calculate fold drops in VE_50_.

### Phenotyping of ACE2 and TMPRSS2 on cell lines

Cell lines (2 × 10^5^ cells) were labelled with phycoerythrin-conjugated TMPRSS2 (Clone: S20014A, BioLegend) and AlexaFluor 647-conjugated ACE2 (Clone: 535919, R&D Systems) for 30 mins on ice in the dark. Cells were washed once with FACS wash (phosphate buffered saline containing 2 mM EDTA and 1% heat-inactivated foetal bovine serum), prior to fixation with 1% paraformaldehyde (final) and acquisition by a BD LSRFortessa^TM^ flow cytometer (BD Biosciences). Flow cytometry analysis was performed using FlowJo analysis software (version 10.8.0, BD Biosciences).

### Detection of catalytic ACE2 in cell supernatants

Soluble ACE2 activity in culture supernatants was measured as previously described ^1^.

### Single resolution of known SARS-CoV-2 entry factors with ACE2 associated solute protein carriers

Single cell RNAseq data from nasal ^57^ and ileum ^58^ were obtained from covid19cellatlas.org, while lung data ^59^ was obtained from GEO (GSE171524). These data were analyzed with Seurat V4.3.0 as described previously ^60^. Their accompanying metadata, which includes information such as sample ID, sample status, and cluster annotations (cell types), were added to Seurat objects using the “AddMetaData” function. Read counts were normalized using SCTransform, before reanalysis with the standard Seurat workflow of “RunPCA,” “FindNeighbours,” “FindClusters,” and “RunUMAP.” Cluster identities were assigned using published cluster annotations and plots were generated with “DimPlot”, “Featureplot” and “DotPlot”, to illustrate expression of *ACE2*, *TMPRSS2*, *SLC6A14*, *SLC6A19* and *SLC6A20*.

### Statistical analysis

Statistical analyses were performed using GraphPad Prism 9 (version 9.1.2, GraphPad software, USA). Sigmoidal dose response curves and interpolated IC50 values were determined using Sigmoidal, 4PL model of regression analysis in GraphPad Prism. For statistical significance, the datasets were initially assessed for Gaussian distribution, based on which further analysis was performed. For datasets that followed normal distribution, unpaired t-test was used to compare two groups. Details of statistical tests used for different data sets have also been provided in figure legends.

---

**Supplementary Figure 1.**
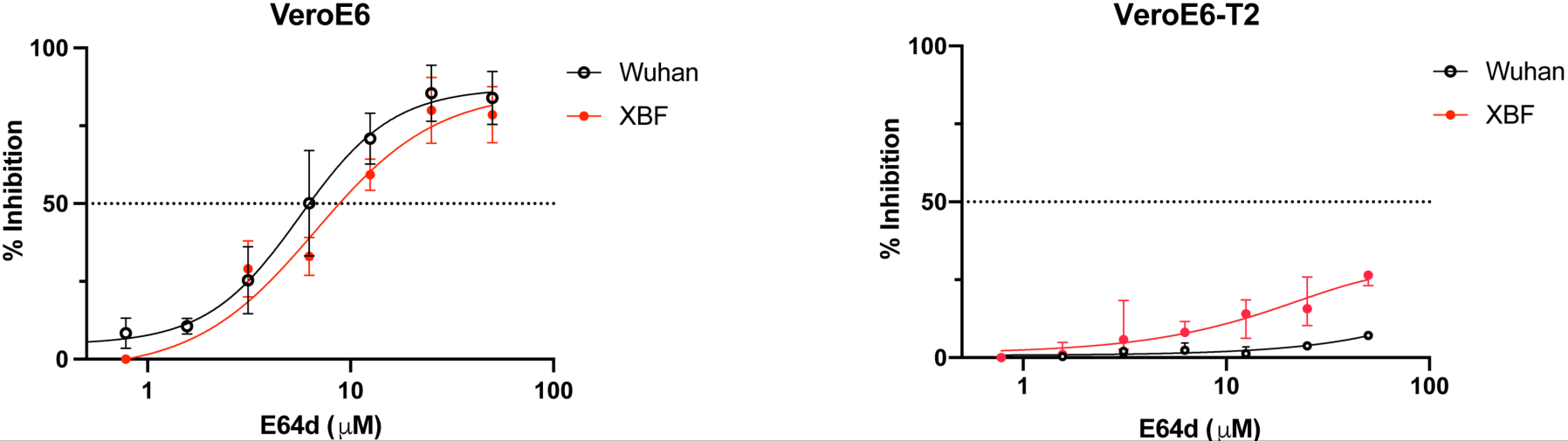
Sensitivity of early and late SARS CoV-2 variants to E64D inhibition in VeroeE6 versus the VeroE6-T2 Cell line A. Unmodified VeroE6s cells were exposed to increasing levels of the Cathepsin L Inhibitor E64D and then were infected with two isolates that represent early (A2.2.) versus contemporary (XBF) variants. B. Lack of inhibition of the pre-Omicron A2.2 versus the contemporary Omicron XBF in the VeroE6-T2 cell line.

**Supplementary Figure 2.**
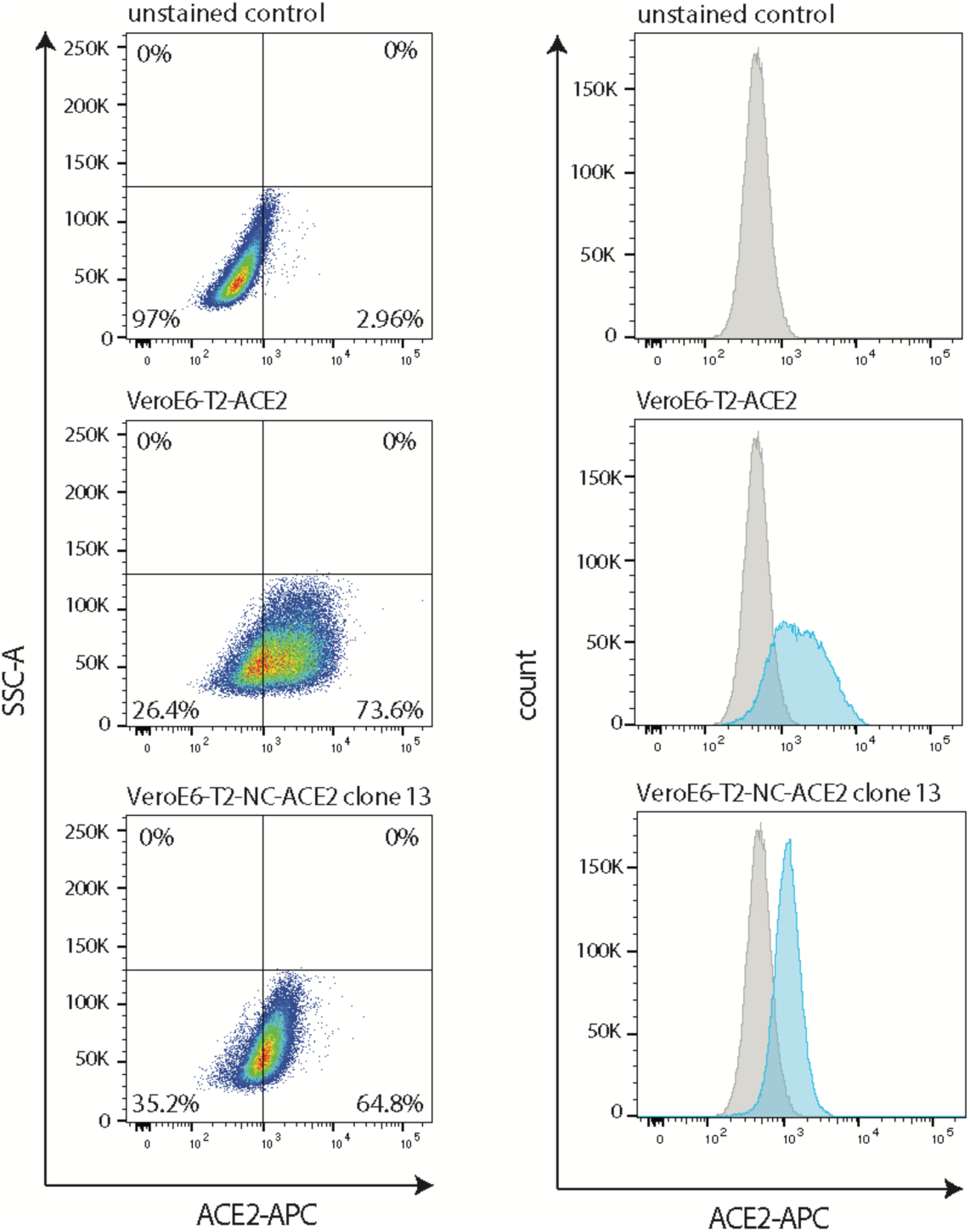
ACE2 surface expression of the VeroE6-T2 cell line with WT ACE2 and TMPRSS2 resistant ACE2 (NC-ACE2). Upper panel: Unstained control. Middle panel: VeroE6-T2-ACE2 Lower panel: VeroE6-T2-NC ACE2 (TMPRSS2 resistant ACE2).

